# Macroscopic dynamics of gene regulatory networks revealed by individual entropy

**DOI:** 10.1101/2021.10.01.462839

**Authors:** Cong Liu, Lijie Hao, Jinzhi Lei

## Abstract

Complex systems are usually high-dimensional with intricate interactions among internal components, and may display complicated dynamics under different conditions. While it is difficult to measure detail dynamics of each component, proper macroscopic description of a complex system is crucial for quantitative studies. In biological systems, each cell is a complex system containing a huge number of molecular components that are interconnected with each other through intricate molecular interaction networks. Here, we consider gene regulatory networks in a cell, and introduce individual entropy as a macroscopic variable to quantify the transcriptional dynamics in response to changes in random perturbations and/or network structures. The proposed individual entropy measures the information entropy of a system at each instant with respect to a basal reference state, and may provide temporal dynamics to characterize switches of system states. Individual entropy provides a method to quantify the stationary macroscopic dynamics of a gene set that is dependent on the gene regulation connections, and can be served as an indicator for the evolution of network structure variation. Moreover, the individual entropy with reference to a preceding state enable us to characterize different dynamic patterns generated from varying network structures. Our results show that the proposed individual entropy can be a valuable macroscopic variable of complex systems in characterizing the transition processes from order to disorder dynamics, and to identify the critical events during the transition process.

## 1 Introduction

Complex systems exist widely in physical, biological, ecological, and social sciences, often compose a large number of strongly interdependent components, and show unanticipated consequences upon small changes in system inputs or structures[1, 2, 3, 4]. From a statistical physical point of view, detail descriptions of each component in a complex system may be entirely intractable (*e.g*., location and velocity of each molecule in a gas), characterizing the macroscopic properties may not only be tractable but also provide us important information of the system (*e.g*., the pressure, temperature, density, compressibility, *etc*.). In biological systems, a single cell is a complex system with a great number of molecules, the molecules connect to each other through molecular interaction networks and their numbers dynamically change over time[5, 6, 7]. Nowadays, advances in a multitude of single-cell sequencing technologies have provide opportunities for direct measurement of molecular details inside a cell[8, 9, 10]. There is an urgent need to develop proper macroscopic variables of a single cell based upon molecule details that can quantify the dynamic process of cell states.

A single cell contains thousands of genes whose expressions are regulated by molecular interplay between DNA, RNA, and proteins, and form a complex system of extremely high dimensional. The signal pathways to regulate the expression process of genes form a gene regulatory network (GRN) that control cellular differentiation and govern the phenotype of an adult somatic cell[11]. Single-cell RNA sequencing technology has made it possible to trace cellular transcriptome at cell level, thereby gives microscopic state of gene transcription in a cell. Biologically, the phenotype of a cell is usually identified by marker genes. Alternatively, low dimensional variables defined by single-cell data may provide more insightful information of a cell that cannot be seen by specific marker genes. Recently, a novel concept of single-cell entropy (scEntropy) was proposed to measure the intrinsic transcriptional state of a cell based on single-cell RNA-seq data[12]. The scEntropy provides a macroscopic variable that can be used to measure the process of early human embryonic development[12], to distinguish malignant cells from normal tissue cells by quantifying the transcriptional order[12], and to describe the dynamics of reprogramming and differentiation of induced pluripotent stem cells[13]. Here, we asked how the data driven concept of scEntropy be extended to model driven complex dynamical systems. Whether the concept of scEntropy is able to quantify the kinetics of transcription state transitions that often show noticeable changes in gene expression patterns? How the concept is incorporated with the complex gene regulatory network dynamics to characterize the gene expression profile?

In this paper, we consider gene regulatory networks modeled with differential equations with random perturbations, propose the concept of individual entropy that extends scEntropy to incorporate the temporal dynamics of complex systems, and investigated how individual entropy can be applied to quantify the transition of system state from stationary to oscillatory dynamics. The original scEntropy was defined as the information entropy of the difference in transcriptions between the cell and a predefined reference cell. In defining the individual entropy, we select proper reference state in accordance with either initial conditions or sliding windows, and consider the system state at each instant as an individual microsystem. The proposed individual entropy provides a method to quantify the stationary macroscopic dynamics of a gene set, and can be served as an indicator for the evolution of network structure variation. Moreover, the individual entropy with proper sliding window based reference enable us to characterize different dynamic patterns generated from varying network structures.

## 2 Method and Model

### 2.1 Individual entropy

The concept of scEntropy was first proposed to measure the order of cellular transcription from single-cell RNA sequencing data with respect to a reference level, larger entropy means lower order in the transcriptions[12]. Consider a gene set of *M* genes, let **r** ∈ ℝ^*M*^ represents the expression vector of reference levels of all genes, and **x** ∈ ℝ^*M*^ the gene expression vector of a cell. scEntropy of the cell **x** with reference to **r**, denoted as *S*(**x**|**r**), is defined as the information entropy of the signal sequence of the difference **y** = **x** − **r**. Explicitly, let

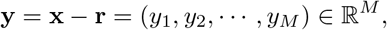

the values {*y*_*i*_} are only informations for the cell **x** (with reference to **r**). While we overlook the order of genes, only the statistical profile of components *y*_*i*_, defined by the probability density function *p*(*y*), is relevant in defining the macroscopic variable. The scEntropy *S*(**x**|**r**) is then defined as

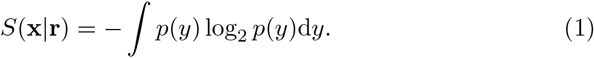

Despite the straightforward and parameter free definition, scEntropy defines an intrinsic transcriptional state of a cell that show results consistent with comment sense observations, *e.g*., increasing entropy along with cell differentiation and larger value entropies for disorder malignant cells[12, 13].

Now, consider dynamics of a complex system with state variables **x**(*t*) = (*x*_1_(*t*), …, *x*_*M*_ (*t*)) ∈ ℝ^*M*^, the dimension *M* is often a large number. Usually, the components *x*_*i*_ are normalized to yield the same dimension values, *e.g*., nondimensionalized variables. Thus, similar to the above definition, let **r**∈ ℝ^*M*^ a reference state, the individual entropy of the system at time *t*, with reference to **r**, is defined as

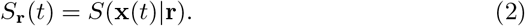

Here, we note that the reference state **r** is a variable for the individual entropy, different choices of the reference state should yield different interpretation of *S*_**r**_(*t*).

When the system dynamics **x** is described by a differential equation (either deterministic or stochastic) with initial condition of form

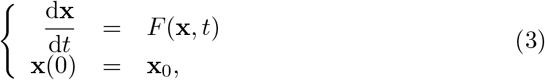

it is straightforward to select the initial state **x**_0_ as the reference state. Hence, we obtain a temporal evolution of the individual entropy

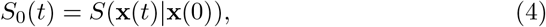

and, obviously, *S*_0_(0) = 0. The entropy *S*_0_(*t*) measures the variance of the order of states **x**(*t*) with respect to the initial condition.

We can also select dynamics reference states as the preceding state with lag time *s*, so that the individual entropy with reference to the preceding states as

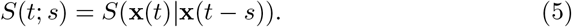

Here, the lag time *s* is the only parameter to define the individual entropy. It is easy to see that *S*(*t*; *s*) is the information entropy for the sequence

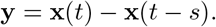

Hence, *S*(*t*; *s*) can be used to measure the changes in the macroscopic state in a time step *s*.

The above definitions *S*_0_(*t*) and *S*(*t*; *s*) are general for any complex systems dynamics. Physically, the individual entropy measures the information of system state **x**(*t*) away from the reference state, and hence provides a macroscopic measurement for the order of system states while we overlook detail interactions in the system. In this paper, we consider specific systems of gene regulatory networks, and investigate how the entropies are applied to study the transition of system dynamics.

### 2.2 Formulation of a gene regulatory network

Now, we considered a system of *M* genes *g*_*i*_ (*i* = 1, 2, …, *M*), and let *x*_*i*_ represents the expression level of the gene *g*_*i*_. The system state is represent by **x** = (*x*_1_, *x*_2_, …, *x*_*M*_). Expression of each gene can be repressed or promoted by other genes, and hence form a complex gene regulatory network. The network structure of regulation relationships is represent by a matrix *V* = (*v*_*ij*_), so that

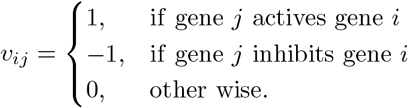

To formulate gene expression dynamics under the regulation network, we need to formulate the overall activation or inhibition effects of all genes to a given gene, which can be biologically complicated with delicate structures[14]. The exact form of mathematical formulation is not important in the current study. Here, we only considered a simple form through Hill type functions.

Consider the gene *g*_*i*_, and assume that the effective promotion of all genes activating *g*_*i*_ expression is represented by the weighted average of all active gene expressions, *i.e*., the effective activation factor *w*_*i*_ defined by

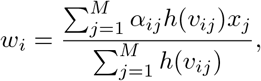

where *α*_*ij*_ is the weighted coefficient of *g*_*j*_ to *g*_*i*_, and *h*(·) is a Heaviside function

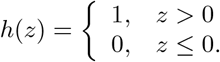

We assumed that the expression rate of *g*_*i*_ depends on the activation factor *w*_*i*_ through an increased Hill type function of form

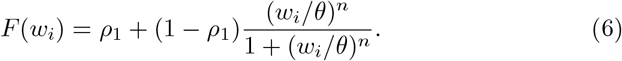

Here *n* represents the Hill coefficient, and *θ* represents the parameter for 50% effective concentration.

Similarly, the effective repression factor *u*_*i*_ for all genes repressing *g*_*i*_ is defined as

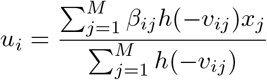

with weighted coefficients *β*_*ij*_. The expression rate of *g*_*i*_ depends on *u*_*i*_ through a decreased Hill type function

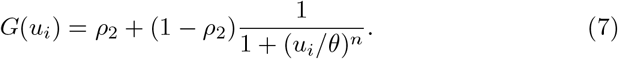

Here, for simplicity, we adopted the same parameters of Hill coefficient and 50% effective concentration as those in the function *F*.

Finally, the dynamic equation of gene expressions were given by differential equations of form

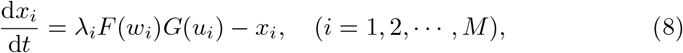

where *λ*_*i*_ is the maximum expression rate of gene *g*_*i*_, and the degradation rates of all genes products are normalized to 1.

Thus, given a gene regulatory network structure *V* = (*v*_*ij*_), the dynamical equations for gene expressions were formulated as

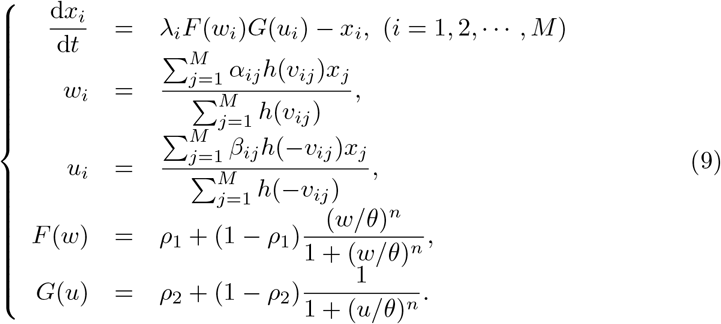

Summary of all notations are given in Table 1.

**Table 1:** Summary of notations.

| Notation | Description |
| --- | --- |
| $x_i$ | expression level of gene $i$ |
| $w_i$ | effective activator factor for gene $g_i$ |
| $u_i$ | effective repression factor for $g_i$ |
| $\alpha_{ij}, \beta_{ij}$ | weighted coefficients |
| $F(w)$ | regulatory activation function |
| $G(u)$ | regulatory inhibition function |
| $\rho_1$ | minimum regulatory activation capacity |
| $\rho_2$ | minimum regulatory inhibition capacity |
| $\lambda_i$ | maximum expression rate of gene $g_i$ |
| $\theta$ | 50% effective concentration of effective regulation factors |
| $n$ | Hill coefficient |
| $v_{ij}$ | regulation of $g_j$ to $g_i$ |
| $\sigma$ | perturbation strength |
| $\tau$ | correlation time of random perturbation |

Next, to consider random perturbations to the gene expression dynamics, we introduced color noise perturbation to model coefficients. Here, we considered external random perturbations to the degradation rates, so that the above equation becomes

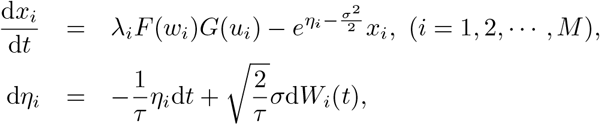

where *W*_*i*_(*t*) is the Weiner process, *τ* is the parameter for the correlation time, and *σ* represents the perturbation strength. Here, *η*_*i*_(*t*) represents an Ornstein-Uhlenbeck process, which satisfies

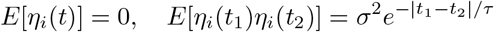

at the stationary state.

In this case, the model equations with random perturbation to degradation rates were formulated as

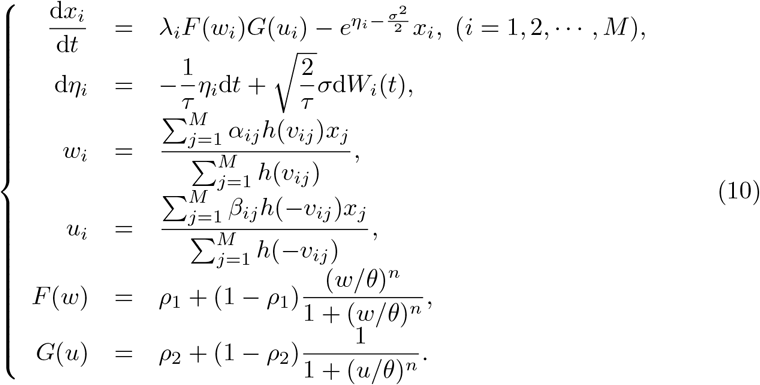

Equations (9) and (10) provide general forms of gene regulatory network dynamics in a cell, and are the main equations considered in the current study.

### 2.3 Network structure

This paper was intended to investigate the method to quantify the macroscopic dynamics of a gene regulatory network, instead of study the exact genetic network in a cell. Hence, we did not consider real signaling networks that are mostly not well understood[15]. We considered specific gene regulatory networks generated from a regular structure involved both promotion and repression interactions. The networks were designed so that system states display transition from order to disorder dynamics, either due to random perturbation or modifications in model structures.

First, we construed a regular network structure *V*_1_ of *M* genes, so that each gene *g*_*i*_ is self-activation and be activated by the preceding gene *g*_*i−* 1_, and is repressed by *g*_*i−* 2_ (Fig. 1a), *i.e*., *v*_*i,i*_ = *v*_*i,i−* 1_ = 1, *v*_*i,i−* 2_ = − 1. Moreover, all genes are located in a circle so that *g*_*M*+*i*_ = *g*_*i*_ for *i >* 0. The structure *V*_1_ is represented as a matrix form of (Fig. 1b)

**Figure 1:**
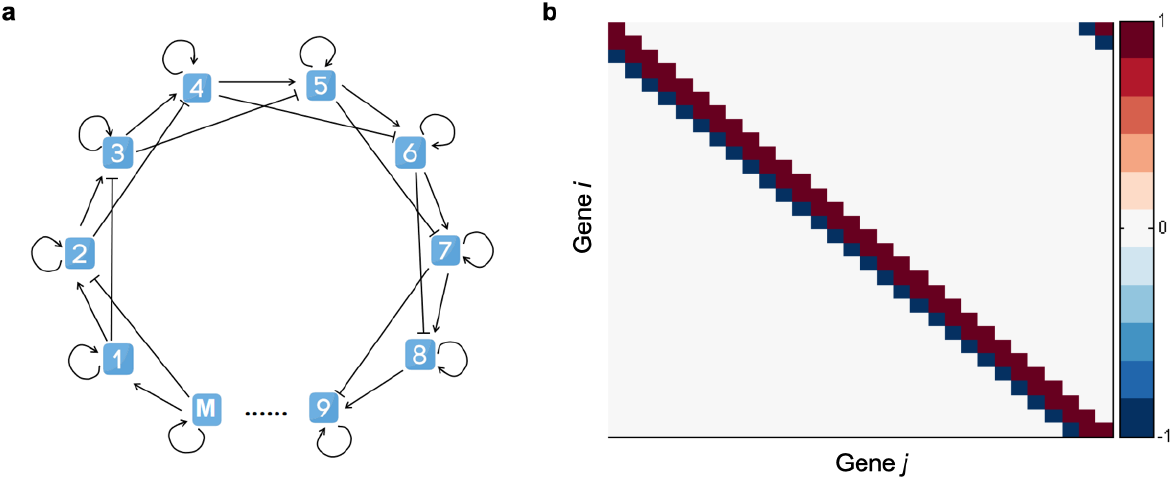
Diagram of the network structure *V*_1_. **a**. Schematic diagram of *M* genes. The blue boxes represent different genes, ‘→’ indicates promotion interaction between genes and ‘⊣’ indicates repression interaction between genes. **b**. Heat map of network structure *V*_1_ (here, *M* = 30). Each color unit indicates the interaction of gene *g*_*j*_ on gene *g*_*i*_.

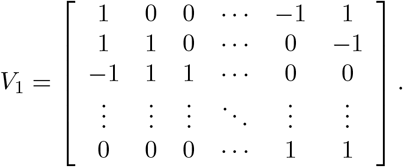

Starting from the regular network *V*_1_, we randomly rewired each edge to obtain an irregular network *V*_2_ so that the system may display disorder dynamics. Biologically, the rewiring process is an analogy of the process of mutation in gene regulations. Transcription networks in a cell show features of a small-world network[16], and the procedure of rewiring edges from a regular network is a standard way to generate a small-world network[17].

## 3 Results

### 3.1 Bifurcation analysis

To identify the parameters used in our study, we performed bifurcation analysis to investigate the possible dynamics of the system with varying parameter values. First, to simplify the bifurcation analysis, we considered the regular regulation network *V*_1_, and applied the symmetric assumption so that

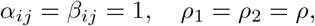

and assumed that the maximum expression rates of all genes are the same

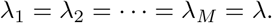

Moreover, we always assumed *θ* = 1 and *n* = 4 in following studies. Therefore, we have only two parameters *λ* and *ρ* in the deterministic equation (9).

First, we considered a simple case with 3 genes, and the model equation (9) becomes

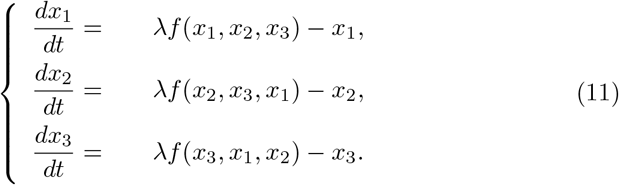

where

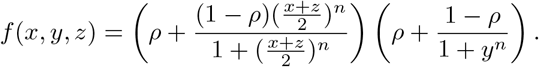

Let *ρ* = 0.1 and applied bifurcation analysis with XPPAUT[18], we obtained a bifurcation diagram shown in Fig. 2a. There are saddle-node bifurcations *λ*_1_(= 2.9956) and *λ*_2_(= 3.4589) so that three steady states exist *λ*_1_ *< λ < λ*_2_ (shadow in Fig. 2a), and the steady state with lower value *x*_1_ is stable. Moreover, there are Hopf bifurcations *λ*_3_(= 2.9994) and *λ*_4_(= 9.1642), so that the steady state with higher value *x*_1_ is unstable when *λ*_3_ *< λ < λ*_4_ and there exists an oscillatory solution.

**Figure 2:**
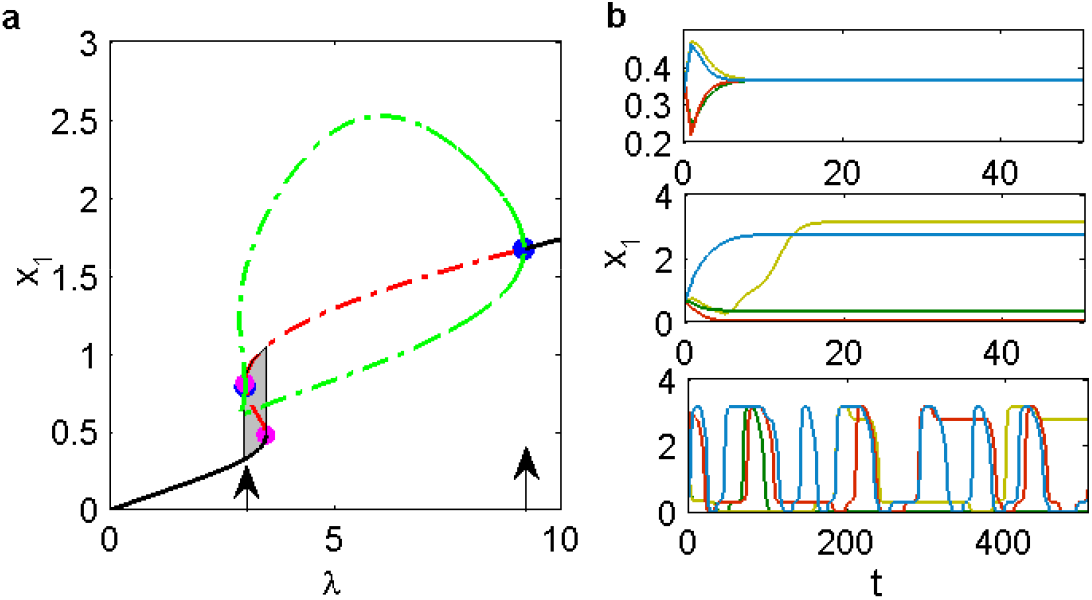
Bifurcation diagram of the deterministic equation. **a**. Bifurcation diagram of the three-dimensional equation (11). The horizontal axis represents the bifurcation parameter *λ*, the vertical axis represents the component *x*_1_. Solid black line represents the stable steady state, red dashed line represents the unstable steady state, green dashed lines represent the upper and lower bound of oscillation solutions generated by Hopf bifurcations, the blue solid dots pointed by the two arrows are Hope bifurcation points, and the solid purple dots are the saddle-node bifurcations. The grey shadow region represents the range of bifurcation parameter *λ* to yield three equilibrium states. Other parameters are *ρ* = 0.1, *n* = 4. **b**. Sample solutions of the model equation (9) (here, *M* = 100) started from different initial conditions. Horizontal axis represents the time *t* and vertical axis represents the component *x*_1_. The top, middle, and bottom panels show solutions started from initial conditions near *x*_*i*_ ≈ 0.371, *x*_*i*_ ≈ 0.641, and *x*_*i*_ ≈ 0.955, respectively. Different colored curves in the graph represent the solution curves starting from different initial values.

Base on the above bifurcation analysis, we take *λ* = 3.2 (*λ*_3_ *< λ < λ*_2_) in this study, so that there is co-existence of stable steady state and oscillatory state. In particularly, while we assume a symmetric steady state *x*_1_ = *x*_2_ = *x*_3_ = *x*^*∗*^, then *x*^*∗*^ satisfies the equation

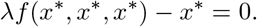

When *λ* = 3.2, there are three solutions *x*^1,*∗*^ ≈ 0.371, *x*^2,*∗*^ ≈ 0.641, and *x*^3,*∗*^ ≈ 0.955, so that (11) has three steady states *E*^*i,∗*^ = (*x*^*i,∗*^, *x*^*i,∗*^, *x*^*i,∗*^), (*i* = 1, 2, 3). Moreover, the state *E*^1,*∗*^ is stable, and the other two states are unstable. Here, we note that the steady states *E*^*i,∗*^ only include those states with equal components, the system may also possess steady states with unequal components. Specifically, when the dimension *M* is large, there are multiple steady states with unequal components.

Based on the above analysis, we set *λ* = 3.2 and considered the situation with a large number of genes (*M* = 100). System dynamics depend on the initial conditions, either converge to stable steady states or oscillatory states (Fig. 2b). Here, we note the existence of multiple steady states, and the system dynamics may sensitively depend on the initial conditions.

### 3.2 Individual entropy with varying initial conditions

From the above analysis, the gene network dynamics show multi-stability, and may display different type dynamics depending on the varying initial conditions. To quantify the dependence of macroscopic dynamics with initial conditions, we took the perturbation strength *σ* = 0.1, and varied the initial conditions over three regions 0.366 *< x*_*i*_ *<* 0.376, 0.636 *< x*_*i*_ *<* 0.646, 0.95 *< x*_*i*_ *<* 0.96, respectively, and solved the equation (10) for dynamic trajectories. The solution dynamics show increasing complexity with varying initial conditions, from homogeneous values, to heterogeneous values, and oscillatory dynamics (Fig. 3a-c, upper panels). Moreover, the distributions of system states for different type dynamics are distinguishable to each other (Fig. 3).

**Figure 3:**
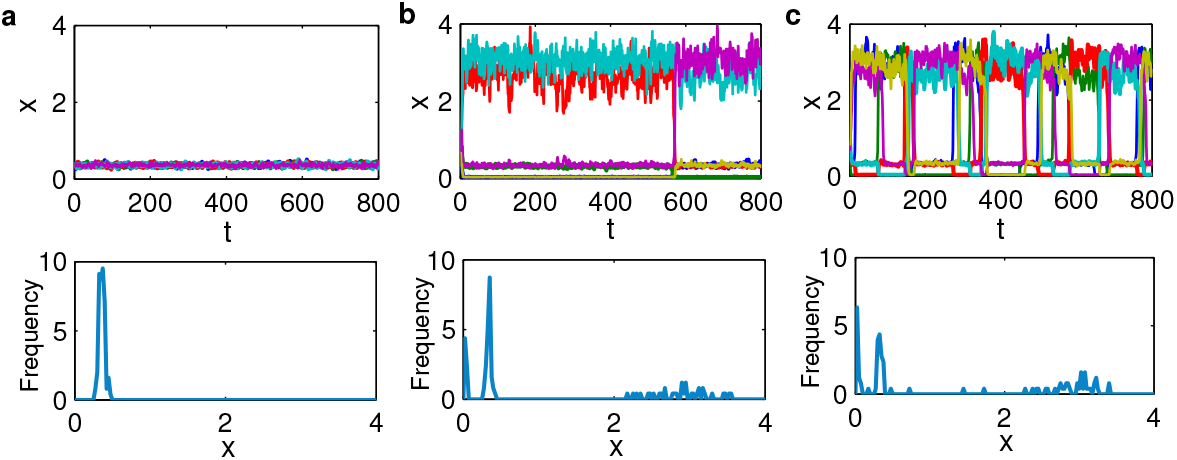
System dynamics under random perturbations. Solution trajectories with different initial conditions, corresponding to (**a**) *x*_*i*_ ≈ 0.371, (**b**) *x*_*i*_ ≈ 0.641, and (**c**) *x*_*i*_ ≈ 0.955, respectively, and noise perturbation with *σ* = 0.1. Upper panels show dynamics of different components (only 5 components are shown with different color curves for clarity), bottom panels show the corresponding distribution of system states {*x*_*i*_(*t*)}_1*≤ i≤ M*_ at time *t* = 800. Here, *M* = 100.

For each initial value regions, we randomly solve 100 independent trajectories, each show different dynamics depending on the initial conditions. We calculated the individual entropy *S*_0_(*t*) for each trajectory, and examine the stationary individual entropy at *t* = 800. The distribution of stationary individual entropies with varying initial conditions is shown in Fig. 4. The distribution of entropies shown well separate for the three region initial conditions, and increase with the increasing complexity of system dynamics. This result indicates that individual entropy can be used to characterize the dynamic complexity of the system with different initial conditions.

**Figure 4:**
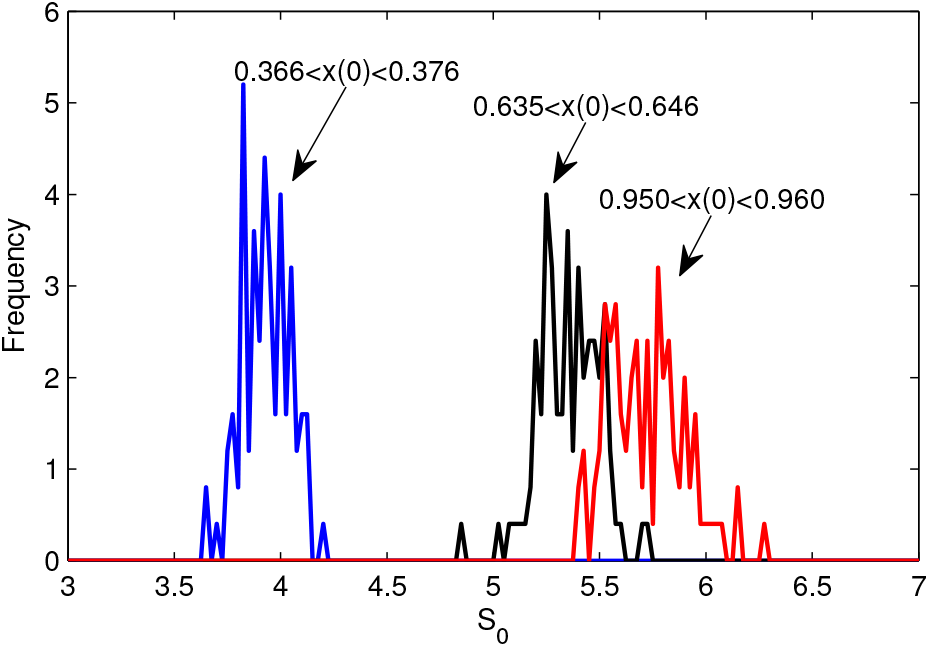
Distribution of individual entropies under different trajectories with different initial values at *t* = 800. Blue, black, and red curves correspond to the initial conditions over three regions 0.366 *< x*_*i*_(0) *<* 0.376, 0.635 *< x*_*i*_(0) *<* 0.646, and 0.950 *< x*_*i*_(0) *<* 0.960, respectively.

### 3.3 Individual entropy to quantify the transition dynamics

To examine how individual entropy can be applied to quantify the transition dynamics, we varied the perturbation strength *σ* so that the system dynamics show transition from low to high complexity. To this end, we initialized the system states 0.366 *< x*_*i*_ *<* 0.376, and set *σ* linearly increase with time *t* as follows

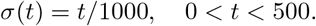

The system dynamics show obviously fluctuation along with the increasing of *σ*, and switches to oscillation dynamics at about *t* = 200 (*σ* = 0.2). To quantify the switches dynamics, we calculated the individual entropy *S*_0_(*t*) with reference to the initial condition. The entropy shows gradually increase from *S*_0_ = 0 at the initial state to a high level of *S*_0_ = 5.8 after the system switches to oscillatory dynamics, akin to the situation with initial condition 0.95 *< x*_*i*_ *<* 0.96 in Fig. 4.

Changes of the individual entropy mainly originate from two sources, the increasing in noise perturbations and the occurring of oscillatory dynamics.

To study how changes in the individual entropy associate with the dynamical switches, we fitted the simulated individual entropy. We note that the simulated data in Fig. 5b presents two phases increases, and each segment increase is in line with a Hill type function. Thus, we fitted the simulated individual entropy with an additive function of two parts as

**Figure 5:**
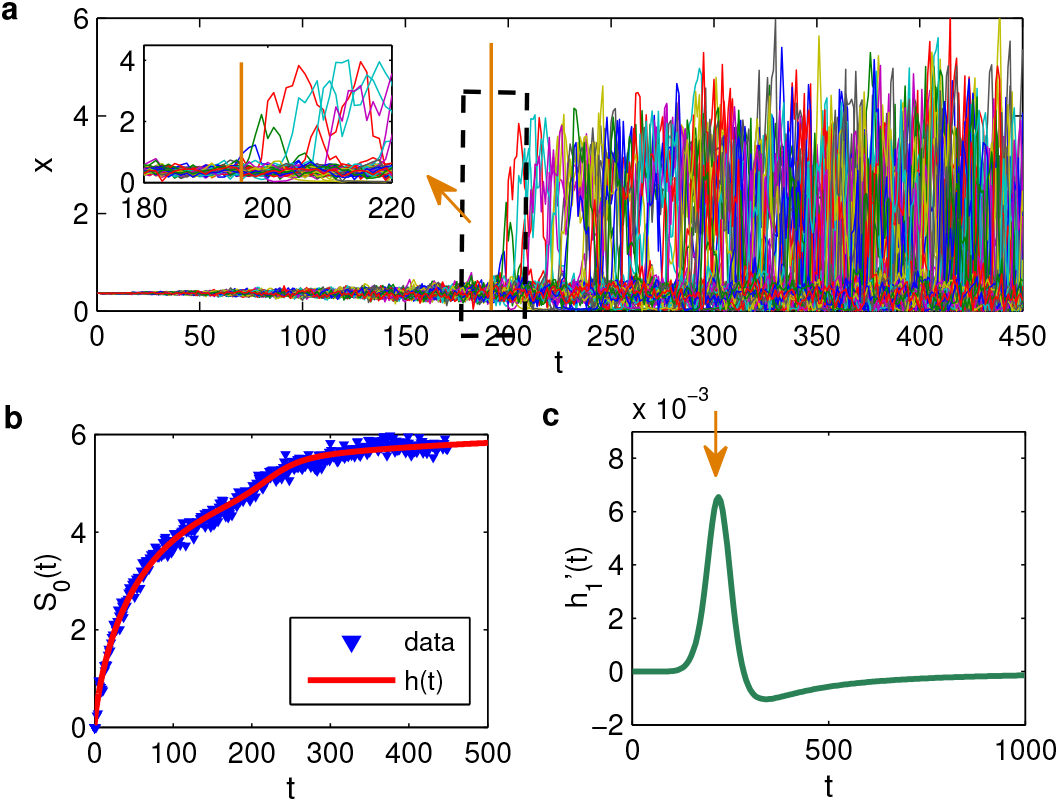
Transition dynamics characterized by the individual entropy. **a**. Evolution of system dynamics with increasing random perturbation strength *σ*. Vertical axis show dynamics of all state variables. Inset show the enlarged of transition dynamics with 180 < *t* < 220. Vertical orange line shows the time of transition that pointed by the arrow in c. **b**. Evolution of individual entropy along the transition dynamics. Blue triangles are obtained from original data, and the red solid line show the fitting function *h*(*t*) = *h*_0_(*t*) + *h*_1_(*t*). **c**. The derivative 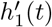. Orange arrow shows the transition time *t* marked with the peak value of 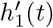.

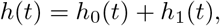

where *h*_0_ and *h*_1_ are Hill type functions defined as

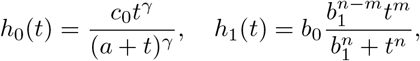

and the parameters *γ* = 0.75, *m* = 10, *n* = 11, *a* = 95.0312, *b*_0_ = 0.622351, *b*_1_ = 230, *c*_0_ = 6.31048. To obtained the parameters, we manually turned the parameters *γ, m, n, b*_1_, and consequently determined other parameters by fitting *h*(*t*) with simulated data. Here, *h*_0_(*t*) represents the changes in the entropy due to random perturbation, and *h*_1_(*t*) represents the additional changes due to the emerge of oscillation dynamics. Fitting of the function *h*(*t*) with simulated *S*_0_(*t*) is shown in Fig. 5b.

Now, the derivative of *h*(*t*), 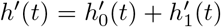, represents the changes of individual entropy *S*_0_(*t*) with *t*. Specifically, the derivative 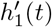 should give the change in entropy due to the emerge of oscillation dynamics, *i.e*., characterization of the transition dynamics. The derivative 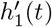 is shown at Fig. 5c, which shows a peak value at *t* = 190 (marked by the arrow in Fig. 5c), corresponding to the occurrence of oscillatory dynamics (Fig. 5a). We further examined the dynamics of each components, the emerge of global oscillatory dynamics starts from oscillations in some components, along which the derivative 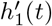 reaches the peak value, and induce the switch of the whole system dynamics. These results suggest that the individual entropy and its derivative can be used to quantify the switches of system state from order to disorder dynamics.

### 3.4 Individual entropy to quantify variations in the network structure

To further investigate how individual entropy is applied to quantify variations in the network structure, we started from the regular network *V*_1_ and randomly rewired the edges to obtain an irregular network *V*_2_ so that system dynamics display oscillations in all components (Fig. 6a). Next, we studied the system dynamics when the network structure randomly change from *V*_1_ to *V*_2_, and examined how individual entropy varies along with changes in the network structure.

**Figure 6:**
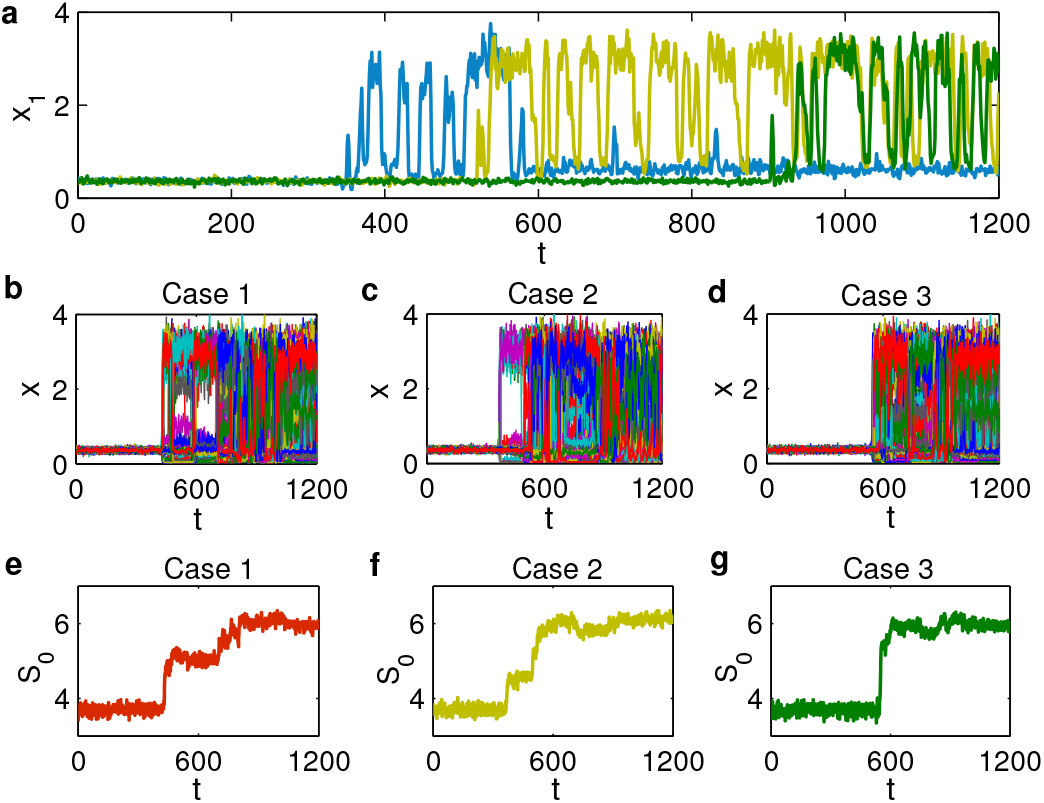
Switches of system dynamics induced by changes in network structures. **a**. System dynamics under network structure *V*_2_. Vertical axis shows the component *x*_1_, and different color lines represent solutions starting from different initial values. **b**-**d**. Different dynamics generated under different pathways from *V*_1_ to *V*_2_. Vertical axis show all state variables *x*_*i*_(*t*). **e**-**g**. Evolution of individual entropies corresponding to different pathways in b-d.

We performed model simulation with network structure randomly changes from *V*_1_ to *V*_2_ according to the scheme

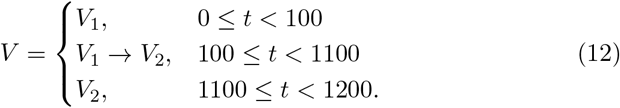

Here, *V*_1_ → *V*_2_ represents a process that randomly change elements in *V*_1_ one-by-one toward *V*_2_ at time *t* = 1100 (detailed in the Appendix). We note that there are multiple pathways from *V*_1_ to *V*_2_, each pathway may yield different dynamics. In performing model simulations, we set the noise perturbation strength *σ* = 0.1, and the initial conditions *x*_*i*_ = 0.371 (*i* = 1, 2, …, *M*). From the above analysis, the system dynamics show small fluctuation around *x*_*i*_(*t*) ≈ 0.371 at early stage when the network structure *V* is close to *V*_1_, and disorder oscillatory dynamics at later stage when *V* is changed toward *V*_2_ (Fig. 6b-d).

Fig. 6b-d show example cases of network structure changes and the switches of system dynamics. Individual entropies *S*_0_(*t*) for each cases are shown in Fig. 6e-g, respectively. From Fig. 6, the system dynamics preserve during the early stage of structure modifications, followed with sudden switch to oscillations in some components, along with a jumping increase in the individual entropy. Moreover, oscillation components may emerge at different time points along the structure modification process, which is in accordance with sudden increases in the individual entropy (Fig. 6e-g). These results suggests that temporal dynamics of the individual entropy *S*_0_(*t*) can effectively reveal the critical moment when there is sudden change in the macroscopic system dynamics.

To further quantify changes of individual entropy along with the process of structure changes, we randomly generated an ensemble of 1000 independent pathways according to the procedure (12), and calculated individual entropies of the system along these pathways. Temporal evolution of the distribution of these individual entropies are shown at Fig. 7. From Fig. 7, despite different dynamics of individual entropy along different pathways of structure modifications (Fig. 6e-g), the entropies are in general increase with changes of the network structure from regular to irregular regulations.

**Figure 7:**
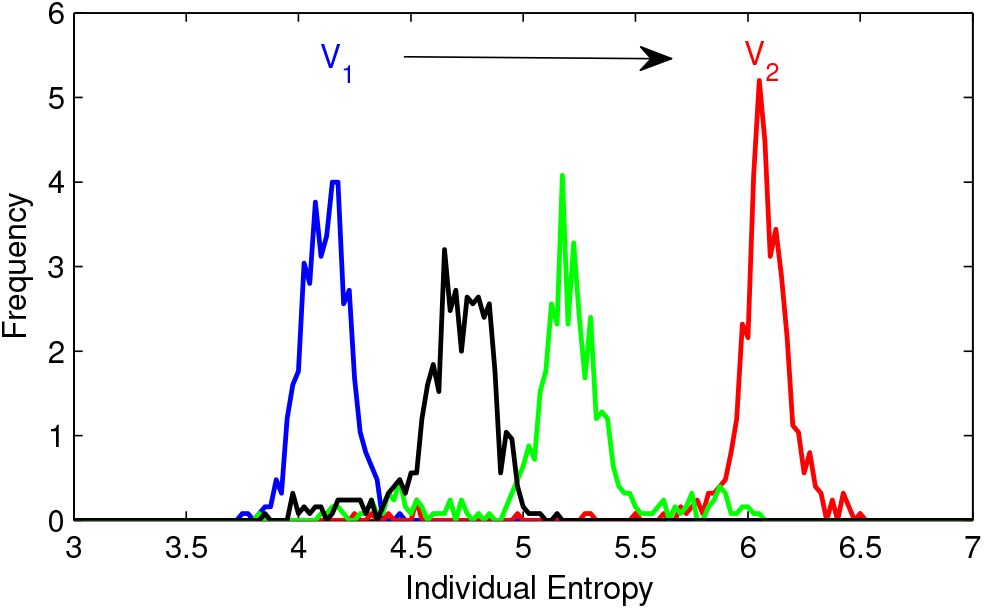
Evolution of the distribution individual entropies along changes of network structure from *V*_1_ to *V*_2_. The fours curves (from left to right) correspond to the time point of *t* = 100, 300, 500, 1100, respectively.

Next, we investigated how individual entropy may quantify the process of reversible change of network structures. To this end, we again start from the structure *V*_1_ and randomly changed the structure to *V*_2_. Next, we randomly recovered the network structure from *V*_2_ to *V*_1_. Thus, the network structure *V* (*t*) is defined as

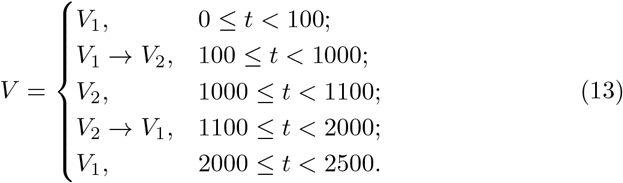

Similar to the previous situation, the system dynamics showed switches between different type dynamics when the structure was changed from *V*_1_ to *V*_2_, from small fluctuations of different components around the same steady states (type I) to random switches between different steady states (type II), and further disorder oscillations (type III) (Fig. 8a). Nevertheless, when the structure recovered from *V*_2_ to *V*_1_, the system dynamics can undergo changes from type III to type II, but not type I dynamics (Fig. 8a). The individual entropy showed increases from *t* = 100 to *t* = 1100 when *V* (*t*) changed from *V*_1_ to *V*_2_, and sudden increases at the moment of switches from type I to type II and from type II to type III dynamics (Fig. 8b). Next, the individual entropy continuously decreases when the structure changed from *V*_2_ to *V*_1_ (*t >* 1100), along with the loss of oscillatory dynamics (Fig. 8b).

**Figure 8:**
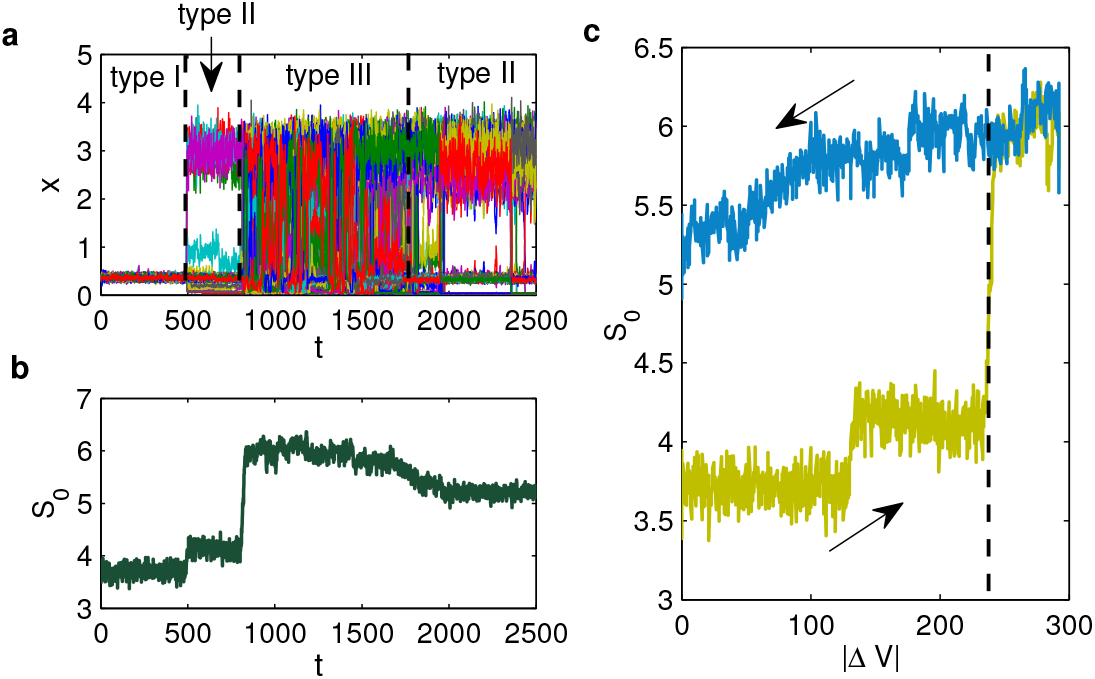
System dynamics in response to reversible change of network structures. **a**. Transition dynamics along with changes of the network structure between *V*_1_ and *V*_2_ follow the equation (13). Vertical axis show all state variables *x*_*i*_(*t*). **b**. Evolution of individual entropy with time *t*. **c**. Dependence of individual entropy with structure variation |Δ*V* |. Arrows show the direction of time increasing.

To measure how the individual entropy may depend on changes in the network structure, we introduced the differences in network structure *V* as

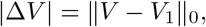

*i.e*., the number of nonzero elements in *V* −*V*_1_. Fig. 8c shows the dependence of individual entropy with |Δ*V* | along the above process. We can see an increase in the entropy when *V* changed from *V*_1_ back to *V*_2_, and a decrease when *V* changed from *V*_1_ to *V*_2_. Interestingly, dependence of the entropy *S*_0_(*t*) with |Δ*V*| is consistent during the reserved process when the difference |Δ*V* | is large (the shadow region in Fig. 8c), and there is a fork bifurcation at a critical value marked by the dashed line in Fig. 8c. There is a sudden increase when |Δ*V* | increase across the critical value, however continuously decrease when |Δ*V* | decrease. These results showed irreversible system dynamics along with changes in network structures, individual entropy can be served as an index to represent the macroscopic system dynamics.

### 3.5 Individual entropy with reference to a preceding state to represent the macroscopic dynamics

In the above transition dynamics (Fig. 6), the system showed type II or III dynamics during the transition of network structures, both with high level individual entropies with reference to initial conditions. To further characterize changes in macroscopic states during the transition process, we considered the individual entropy *S*(*t*; *s*) along the transition dynamics with reference to the preceding state with lag time *s*.

In Fig. 9, we show transition dynamics along with changes of the network structure between *V*_1_ and *V*_2_ follow the equations (12) (Fig. 9a) or (13) (Fig. 9b). Here, to be more focus at the dynamical features of heterogeneity in different components, we omitted the noise perturbation and hence *σ* = 0; moreover, maximum expression rates (*λ*_*i*_) are assumed to be different for each genes, and take values randomly over 3.2 ± 1.0.

**Figure 9:**
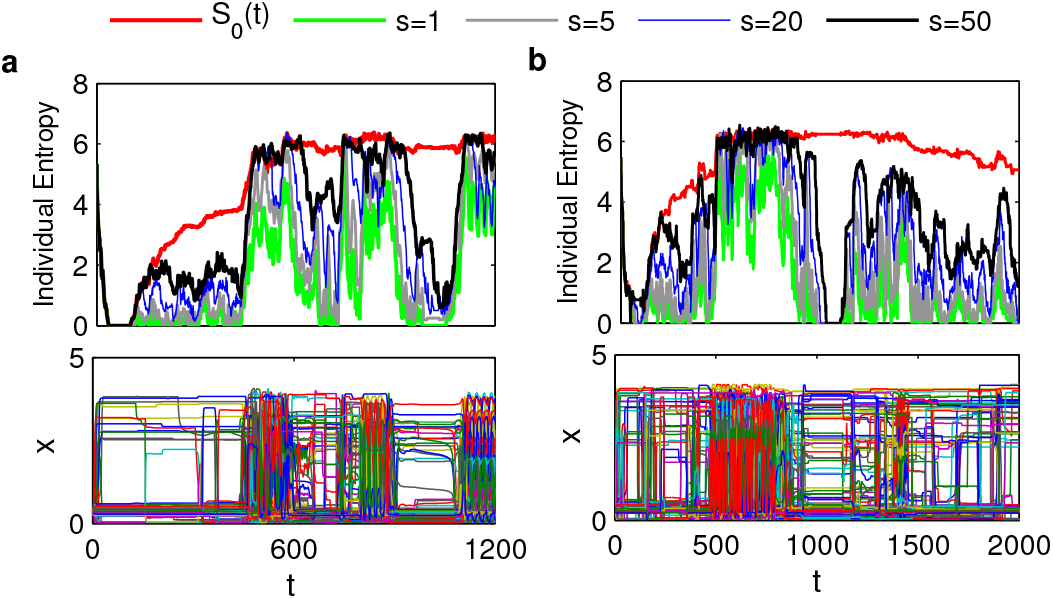
Description of individual entropy on the transition of dynamical patterns. **a**. The network structure changed from *V*_1_ to *V*_2_ according to (12). **b**. The network structure changed between *V*_1_ and *V*_2_ according to (13). The upper panels show plots of individual entropy over time, different color lines indicate different selection of reference states, either with reference to the initial state (*S*_0_(*t*)), or preceding state with different lag time *s* (*S*(*t*; *s*)); bottom panels showed system dynamics over time. Here, the expression rate *λ*_*i*_ ∈ [2.2, 4.2], and the noise strength *σ* = 0.

Dynamical patterns during the transition process are shown in bottom panels in Fig. 9. To quantify the dynamical patterns, we calculated the individual entropy along process with reference to the initial state (*S*_0_(*t*)), and the preceding states with lag time *s* (*S*(*t*; *s*)). When referred to the initial state, the individual entropy show continuously increase during the transition process, and cannot distinguish different dynamical patterns (Fig. 9, red lines in upper panels). Nevertheless, when we referred to preceding states, the individual entropy showed up and down dynamics in accordance with changes in dynamical patterns of system states (Fig. 9). Moreover, we varied the lag time from *s* = 1 to *s* = 50, and the entropy *S*(*t*; *s*) was insensitive with *s*. These results suggest that individual entropy with reference to preceding state provides a reasonable quantity measurement for the dynamical patterns during the transition dynamics.

### 3.6 Comparison with other type entropies

Now, we recall our definition, the proposed individual entropy is the information entropy of the probability function for the signal sequence of the difference **y** = **x**−**r**, denoted as *S*(**x**|**r**). Alternatively, we can directly calculate the information entropy of probability function for the signal sequence of **x**, denoted as *S*(**x**); or, given the reference state, the relative entropy (Kullback-Leibler divergence) of the probability function for the sequence **x** with respect to that for the reference sequence **r**, denoted as *KL*(**x, r**).

To compare how different type definitions may be applied to quantify the system dynamics, we considered again the transition processes in Fig. 6b and Fig. 9a. We set the reference state **r** as the initial condition, and calculated the entropies *S*(**x**(*t*) |**r**), *S*(**x**(*t*)), and *KL*(**x**(*t*), **r**) respectively. The results of various types entropies are shown in Fig. 10.

**Figure 10:**
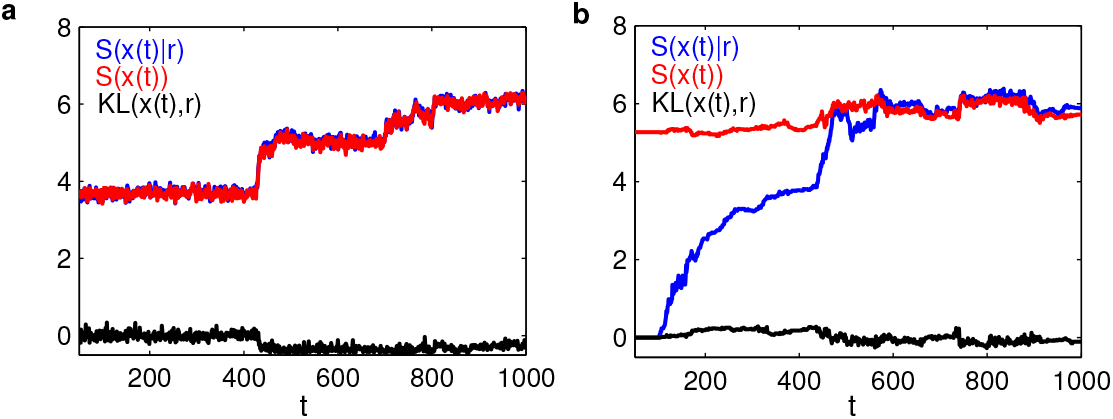
Comparison between different type entropies for the transition processes in (a) Fig. 6b and (b) Fig. 9a, respectively. The horizontal axis is time *t*, the vertical axis represents various types of entropy, and different colors represent the different entropy changing over time. Blue represents individual entropy, red represents absolute entropy, and black represents KL divergence.

In the dynamics shown by Fig. 6b, all components in the reference state **r** take the same value, the sequence **y** = **x**−**r** is simply a translation of **x**, and hence the individual entropy *S*(**x**|**r**) is the same as the information entropy *S*(**x**), both of them are able to characterize the sudden change in dynamical patterns (Fig. 10a). Nevertheless, the Kullback-Leibler divergence *KL*(**x, r**) fail to quantify obvious changes in the system dynamics (Fig. 10a).

In the dynamics shown by Fig.9a, different components in the reference state **r** are different from each other, and hence the individual entropy *S*(**x**|**r**) is different from *S*(**x**). From Fig. 10b, *S*(**x**|**r**) shows more significant difference for different pattern dynamics, which suggest better characterization of macroscopic dynamics than the traditional information entropy *S*(**x**). Moreover, the Kullback-Leibler divergence fail to quantify changes in the system dynamics in this situation (Fig. 10b).

## 4 Conclusions

In this paper, we proposed a concept of individual entropy as a macroscopic measurement of the complexity for complex system dynamics. Individual entropy was defined following a novel concept of single-cell entropy that was originally proposed to measure the intrinsic transcriptional state of a cell based single-cell RNA-seq data[12]. Consider dynamics of a complex system with state variables **x**(*t*), individual entropy of the system state at any time *t*, with reference to a state **r**, is defined as the information entropy of the signal sequences of the difference **y**(*t*) = **x**(*t*) −**r**. The proposed individual entropy provides a macroscopic measurement for the order of system states while we overlook detail interactions in a complex system. This is certainly important in biological systems that detail molecular interactions are mostly unclear.

In our study, we considered dynamical systems from a gene regulatory network as an example to investigate how the concept of individual entropy is applied to quantify the transition dynamics. Results showed that critical events of sudden changes in system dynamics due to random perturbation can be described by variations in the individual entropy (Fig. 5). Moreover, different type dynamics during the transition process induced by network structure alterations can be described by sudden changes in the individual entropy (Figs. 6-8).

In defining the individual entropy, the reference state provides a degree of freedom for the baseline state with the minimum entropy of zero. Different selection of the reference state may yield different interpretation of the results. In our study, we found that the individual entropy with reference to the preceding state with a lag time is able to describe changes in dynamical patterns during state transition induced by changes in the network structure. These results showed that the proposed individual entropy with proper selection of reference states can be a practical variable to quantify complex gene regulation dynamics.

To conclude, main outcomes of the present study concerns the macroscopic dynamics of complex systems. We proposed a concept of individual entropy as an effective indicator to describe the transition dynamics of system states. The current study was performed based on an artificial gene regulatory network. However, the concept can easily be extended to dynamics related with general genetic networks, and hence can guide the studies of cell state transition in various biological processes. Moreover, the concept of individual entropy can also be applied to other complex dynamics systems, and provides a variable quantity for complex systems without enough knowledge in the detail informations of the system.

## Acknowledgments

This work was funded by National Natural Science Foundation of China (NSFC 11831015).

## Appendix: Random changes of network structure

To simulate the random changes of network structure *V*_1_ → *V*_2_ in (12), we let

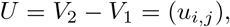

and denote Σ for all nonzero elements *u*_*i,j*_. Assume that Σ contains totally *m* nonzero elements, we divide the time interval *t* ∈ (100, 1100) into *m* time points

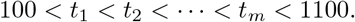

Next, let

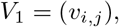

and do the following iteration:
**for** *k* from 1 to *m* **do**

- randomly select one element in Σ, said *u*_*i,j*_;
- let *v*_*i,j*_ = *v*_*i,j*_ + *u*_*i,j*_;
- remove *u*_*i,j*_ from Σ.

This process randomly changes the network structure from *V*_1_ to *V*_2_.

## References

[1] Sunny Y. Auyang. Foundations of Complex-system Theories: in Economics, Evolutionary Biology, and Statistical Physics. Compridge University Press, 1998.

[2] G. Nicolis, I. Prigogine, and P. Carruthers. Exploring complexity: An introduction. Physics Today, 43(10):96–97, 1990.

[3] Avi Ma’ayan. Complex systems biology. J R Soc Interface, 14:20170391, 2017.

[4] Alexander F. Siegenfeld and Yaneer Bar-Yam. An introduction to complex systems science and its applications. Complexity, 2020:6105872, 2020.

[5] Volker Bergen, Marius Lange, Stefan Peidli, F Alexander Wolf, and Fabian J Theis. Generalizing RNA velocity to transient cell states through dynamical modeling. Nat Biotechnol, 38(12):1408–1414, 2020.

[6] BJ Strober, R Elorbany, K Rhodes, N Krishnan, K Tayeb, A Battle, and Y Gilad. Dynamic genetic regulation of gene expression during cellular differentiation. Science, 364(6):1287–1290, 2019.

[7] Gene-Wei Li and X Sunney Xie. Central dogma at the single-molecule level in living cells. Nature, 475(7356):308–315, 2011.

[8] Fuchou Tang, Catalin Barbacioru, Yangzhou Wang, Ellen Nordman, Clarence Lee, et al. mRNA-Seq whole-transcriptome analysis of a single cell. Nat Methods, 6(5):377–382, 2009.

[9] Xianwen Ren, Boxi Kang, and Zemin Zhang. Understanding tumor ecosystems by single-cell sequencing: promises and limitations. Genome Biol, 19(1):211, 2018.

[10] X Sunney Xie, Paul J Choi, Gene-Wei Li, Nam Ki Lee, and Giuseppe Lia. Single-molecule approach to molecular biology in living bacterial cells. Ann Rev Biophys, 37(1):417–444, 2008.

[11] Aditya Pratapa, Amogh P Jalihal, Jeffrey N Law, Aditya Bharadwaj, and TM Murali. Benchmarking algorithms for gene regulatory network inference from single-cell transcriptomic data. Nat Methods, 17(2):147–154, 2020.

[12] J. Liu, Y. Song, and J. Lei. Single-cell entropy to quantify the cellular order parameter from single-cell rna-seq data. Biophys Rev Lett, 15(1):1–15, 2020.

[13] Y. Ye, Z. Yang, M. Zhu, and J. Lei. Using single-cell entropy to describe the dynamics of reprogramming and differentiation of induced pluripotent stem cells. Int J Mod Phys B, 34(30):2050288, 2020.

[14] G. Karlebach and R. Shamir. Modelling and analysis of gene regulatory networks. Nat Rev Mol Cell Biol, 9(10):770–80, 2008.

[15] Xiao-Tai Huang, Yuan Zhu, Lai Hang Leanne Chan, Zhongying Zhao, and Hong Yan. Inference of cellular level signaling networks using single-cell gene expression data in Caenorhabditis elegans reveals mechanisms of cell fate specification. Bioinformatics, 33(10):1528–1535, 2017.

[16] Virginia E Glazier and Damian J Krysan. Transcription factor network efficiency in the regulation of Candida albicans biofilms: it is a small world. Curr Genet, 64:883–888, 2018.

[17] Duncan J. Watts and Steven H. Strogatz. Collective dynamics of ‘smallworld’ networks. Nature, 393:440–442, 1998.

[18] Bard Ermentrout. Simulating, Analyzing, and Animating Dynamical Systems: A Guide to XPPAUT for Researchers and Students. SIAM, 2002.

[19] R. A. Meyers. Encyclopedia of Complexity and Systems Science. Encyclopedia of Complexity and Systems Science, 2009.

[20] H. E. Stanley and G. Ahlers. Introduction to phase transitions and critical phenomena. Phys Today, 26(1):71–72, 1973.

[21] Albert-László Barabási and Réka Albert. Emergence of scaling in random networks. Science, 286:509–512, 1999.

[22] L. Glass and M. C. Mackey. From Clocks to Chaos. Princeton University Press, Princeton NJ, 1988.

[23] P. E. Hardin, J. C. Hall, and M. Rosbash. Feedback of the drosophila period gene product on circadian cycling of its messenger rna levels. Nature, 343(6258):536–540, 1990.

[24] Renato E. Mirollo and Steven H. Strogatz. Synchronization of pulse-coupled biological oscillators. SIAM J Appl Math, 50(6):1645–1662, 1990.

[25] C. S. Peskin. Mathematical aspects of heart physiology. York University, 1975.

[26] Y. Kuramoto. Chemical oscillations, waves, and turbulence. Springer-Verlag, 1984.

[27] Zoltán N. Oltval and Albert-László Barabási. Life’s complexity pyramid. Science, 298(5594):763–764, 2002.

[28] T. Schlitt and A. Brazma. Modelling gene networks at different organisational levels. Febs Lett, 579(8), 2005.

[29] J. Hasty, D. Mcmillen, F Isaacs, and J. J. Collins. Computational studies of gene regulatory networks: in numero molecular biology. Nat Rev Genet, 2(4):268–279, 2001.

[30] Mads Kærn, Timothy C. Elston, William J. Blake, and James J. Collins. Stochasticity in gene expression: from theories to phenotypes. Nat Rev Genet, 6(6):451–64, 2005.

